# A hybrid Embden–Meyerhof–Parnas pathway provides a synthetic link between sugar and phosphate metabolism

**DOI:** 10.1101/2020.06.11.147082

**Authors:** Yoojin Lee, Hye Jeong Cho, Han Min Woo

## Abstract

The fundamental Embden–Meyerhoff–Paranas (EMP) pathway for sugar catabolism, anabolism, and energy metabolism has been reconstituted with non-oxidative glycolysis (NOG). Although carbon conservation was achieved via NOG, the energy metabolism was significantly limited. Herein, we showed the construction of a hybrid EMP that replaced the first phase of the EMP in *Corynebacterium glutamicum* with NOG and revealed a metabolic link of carbon and phosphorus metabolism. In accordance with synthetic glucose kinase activity and phosphoketolase on the hybrid EMP, cell growth was completely recovered in the *C. glutamicum pfkA* mutant strain where the first phase of EMP was eliminated. Notably, we have revealed a phosphate-replenishing pathway that involved trehalose biosynthesis for the generation of inorganic phosphate (P_*i*_) sources in the hybrid EMP when external P_*i*_ supply was limited. Thus, the re-designed hybrid EMP pathway with balanced carbon and phosphorus states provides an efficient microbial platform for biochemical production.

---

The Embden–Meyerhoff–Paranas (EMP) pathway is a fundamental pathway for sugar metabolism in the cell and it is challenging to design a synthetic pathway based on it with reduced CO_2_ loss or conversion of CO_2_^1–5^. In the EMP pathway, an initial investment of ATP during the preparatory phase is necessary for the generation of a total of 2 ATP per glucose molecule at the pay-off phase. Thus, inorganic phosphate (P_*i*_) must be supplied for the EMP pathway in the medium and glucose can be metabolized to supply key metabolites such as pyruvate, C3; acetyl-CoA, and C2. A total loss of 2 CO_2_ molecules per glucose is inevitable during EMP to synthesize acetyl-CoA from pyruvate by the pyruvate dehydrogenase complex. If the CO_2_ wastage is minimized, the theoretical yield can be increased in terms of carbon balance. In nature, a pyruvate by-pass has evolved to conserve the carbon molecules for the pentose sugars by generating C3 and C2 metabolites via the action of a series of enzymes such as sugar kinases, phosphorylase and phosphoketolase (Pkt)^6^. On the other hand, phosphorus is a major nutrient required for cellular processes such as sugar metabolism, which is coupled with phosphorus metabolism (mainly P_*i*_ in the minimal medium) as cells need to dynamically adapt to environmental changes^7,8^. Thus, it is necessary to decipher the pathways for both carbon and phosphorus metabolism while designing a synthetic sugar metabolic pathway to ensure balanced cellular fitness.

Recently, carbon conservation in sugar metabolism has been successfully achieved using synthetic design with Pkt and a carbon rearrangement cycle to generate 3 acetyl-CoA per fructose 6-phosphate (F6P) molecule^9^. Non-oxidative glycolysis (NOG) has provided a biotechnological basis for increasing the theoretical yield by reducing CO_2_ evolution. To fulfill carbon conservation in *Escherichia coli*, adaptive evolution has been performed in a minimal glucose medium after rational metabolic engineering^10^, which results in slower growth of the NOG strains than their parental strain. This could be due to insufficient reducing equivalents generated via NOG. In addition, a bifid shunt, which also involves Pkt, has been introduced in *E. coli* under the same biotechnological principle of NOG^11^. The EP-bifido pathway has enabled the increase in production of acetyl-CoA from acetyl phosphate (AcP) and has been applied to enhance the levels of acetyl-CoA–derived chemicals. Although CO_2_ production was reduced in the EP-bifido pathway, glucose utilization was incomplete in the engineered *E. coli* due to possible metabolic limitations. However, studies have not investigated the metabolic balance between carbon and phosphorus metabolism in the NOG or the EP-bifido pathways, which might be crucial for heterologous Pkt activity. P_*i*_ balance is crucial for Pkt activity in both NOG and Ep-bifido and hence, P_*i*_ availability might affect the activity of phosphoacetyltransferase (Pta) as there is a decrease in the net P_*i*_ in the synthetic pathways (**Fig. 1**). Unless supplemented, cellular metabolism must self-regulate in order to replenish the internal P_*i*_ sources although ATP hydrolysis provides the P_*i*_ in the limited ATP pools.

**Figure 1.**
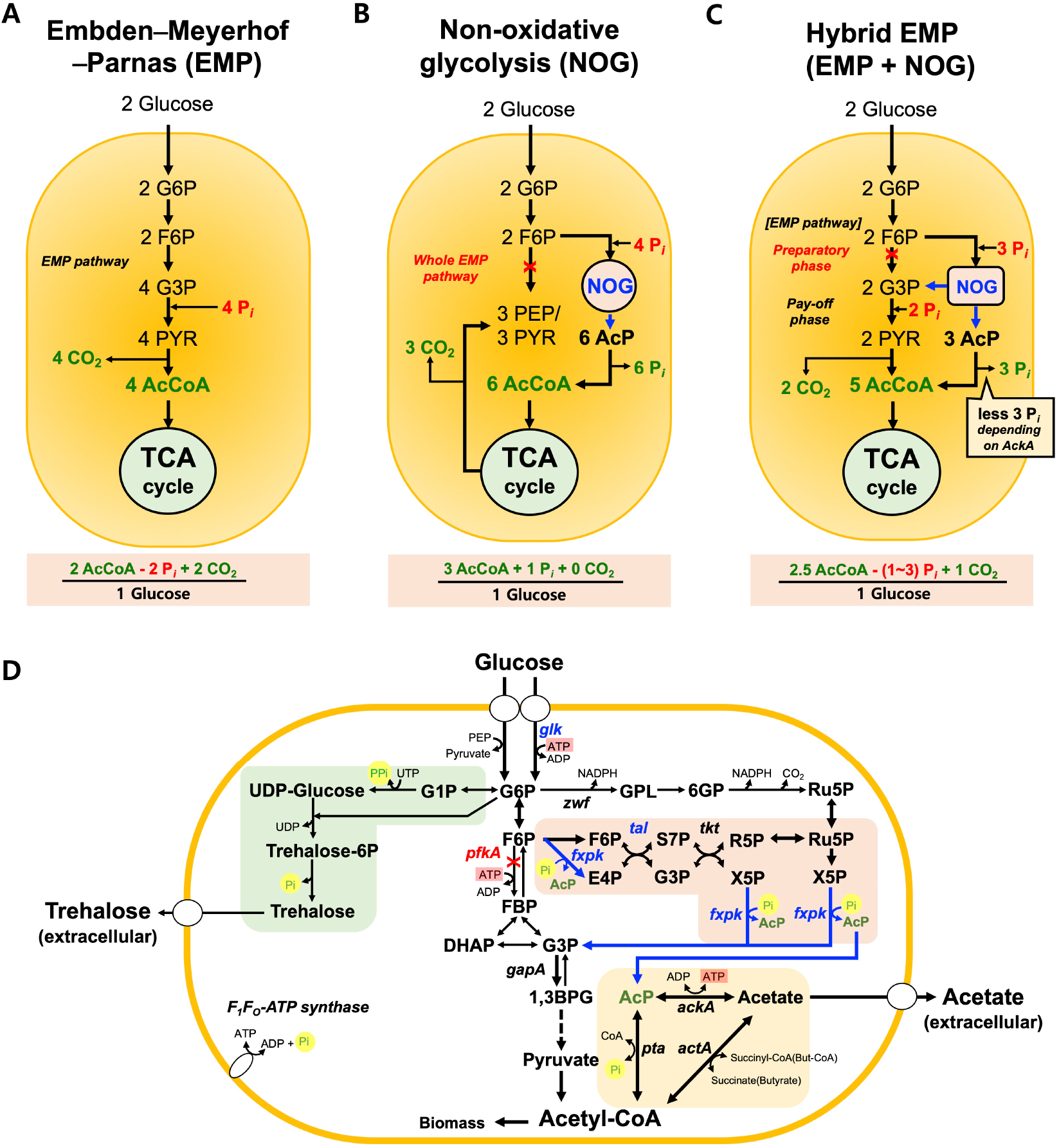
Structure of EMP pathway, NOG, and hybrid EMP pathway. (**A**) a simplified scheme of EMP (or glycolysis) (**B**) a simplified scheme of non-oxidative glycolysis^9,10^ (**C**) a simplified hybrid EMP where the preparatory phase was replaced by the NOG pathway. (**D**) Metabolic networks in the engineered *C. glutamicum* with the hybrid EMP. The red letter and blue letter represent gene deleted or overexpressed, respectively. The black arrow and blue arrow represent native pathway and synthetic pathway, respectively. The green highlighted path indicates trehalose biosynthesis pathway via UDP-glucose. The red highlighted path indicates the NOG pathway. The yellow highlighted path indicates the flexible AcP-acetate pathway. Abbreviations: EMP, Embden–Meyerhof–Parnas; NOG, non-oxidative glycolysis; G6P, glucose 6-phopshate; G1P, glucose 1-phosphate; Trehalose-6P, Trehalose 6-phosphate; F6P, fructose 6-phosphate; PGL, 6-phosphoglucono 1,5,-lactone; 6PG, 6-phosphogluconate; Ru5P, ribulose-5-phosphate; S7P, sedoheptulose 7-phosphate; E4P, erythrose-4-phosphate; X5P, xylulose-5-phosphate; R5P, ribose-5-phosphate; G3P, glyceraldehyde 3-phosphate; DHAP, dihydroxyacetone; 1,3BPG, 1,3-bisphosphate glycerate; PEP, phosphoenolpyruvate; Pyr, Pyruvate; AcP, acetyl phosphate; AcCoA, acetyl-CoA; *glk*, glucokinase; *pfkA*, phosphoglucokinase; gapA, glyceraldehyde 3-phosphate dehydrognease; *zwf*, glucose-6-phosphate 1-dehydrogenase; *tkt*, transketolase; *tal* transaldolase; *fxpk*, bifunctional phosphoketolase; *pta*, phosphotransacetylase*; ackA*, acetate kinase; *actA*, succinyl-CoA:acetate CoA-transferase.

To overcome the aforementioned limitations and to address the questions, we report a metabolic design involving the construction of hybrid EMP to provide sufficient ATP and NADH production for cellular fitness, and to reduce CO_2_ and increase acetyl-CoA for biotechnological applications. To completely understand the glucose catabolism, we examined the fate of AcP as an output of NOG *in vivo* by constructing the mutants at the AcP-AcCoA node with respect to P_*i*_. Notably, we found a metabolic link between the glucose catabolism in the hybrid EMP pathway and the trehalose biosynthesis pathway regulated by the phosphate availability. Thus, cells metabolize glucose in the hybrid EMP pathway to provide elevated acetyl-CoA levels and reduce CO_2_ production. This pathway might be significantly beneficial for glucose utilization without compromising the reducing equivalents along with balanced phosphate metabolism.

## Results

### Designing the hybrid EMP pathway

We designed a synthetic NOG pathway to preserve the carbon molecules during sugar metabolism without loss of CO_2_ and the enzymatic reactions involved in NOG from F6P to AcP have been shown in the *in vitro* carbon conservation cycle (**Fig. 1**)^9^. However, the synthetic NOG strain, in which the native EMP pathway was completely blocked by deleting the key glycolytic genes including *gapA* (glyceraldehyde-3-phosphate dehydrogenase, GapA) and *pfkA* (phosphofructokinase, PfkA), was not able to grow in the minimal glucose medium. Further, the evolution of the engineered *E. coli* led to aerobic cell growth in the minimal glucose medium. However, NOG-evolved strains showed much slower growth rates in the minimal glucose medium than that of the WT due weakened C3 synthesis^10^. Therefore, we designed a hybrid EMP pathway consisting of three phases: (i) an NOG pathway phase that converts F6P to AcP (**eq. 1–1**), (ii) a pay-off phase that generates ATP and NADH by converting G3P to AcCoA (**eq. 2**), and (iii) an AcP–acetyl-CoA node that converts AcP to acetyl-CoA via with or without acetate biosynthesis (**eq. 3**) (**Fig. 2A**).

**Figure 2.**
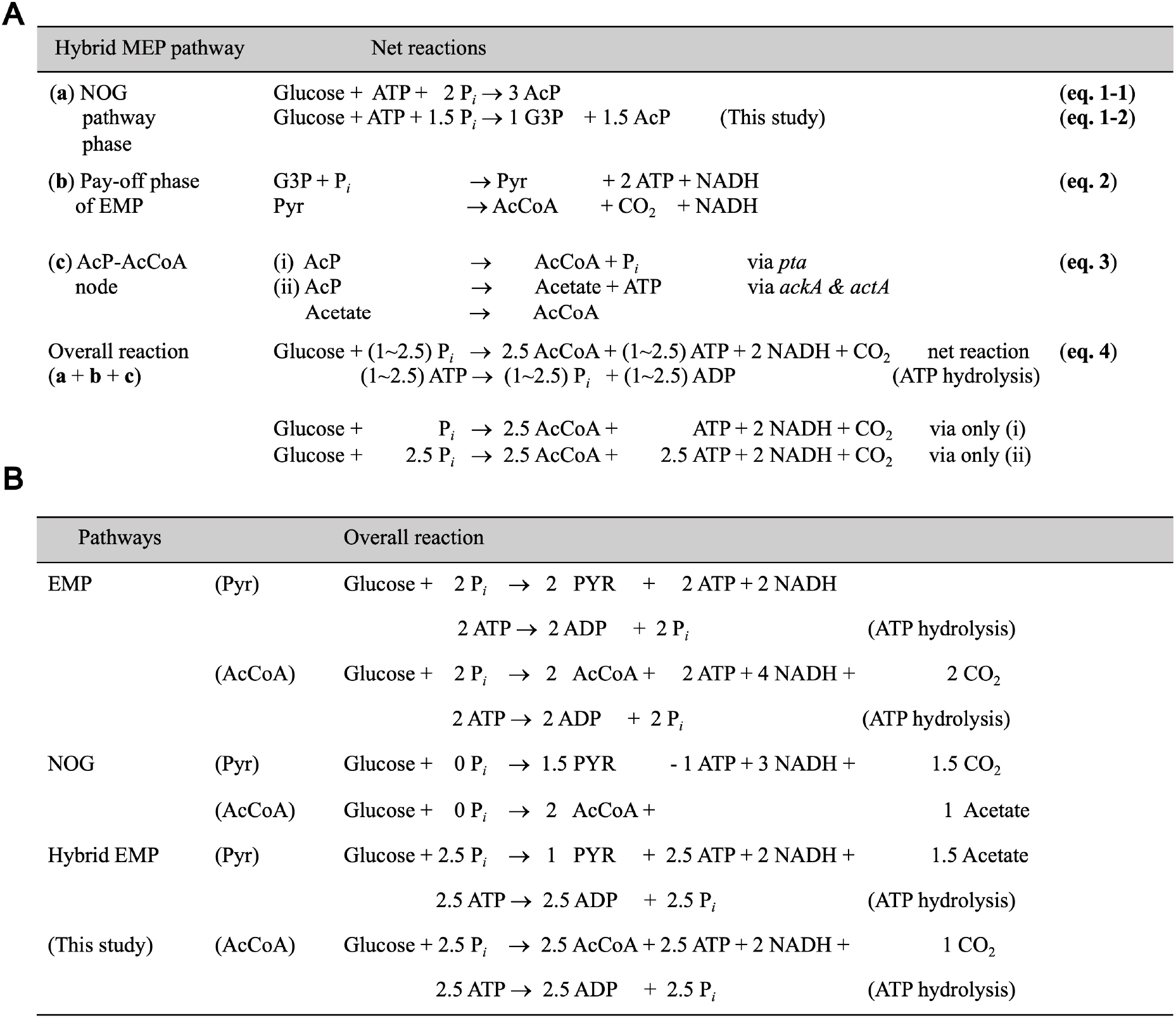
Scheme of the overall reactions for hybrid EMP. (**A**) The hybrid EMP pathway was designed three phases (**a**) NOG pathway, (**b**) the pay-off phase of EMP, (**c**) AcP-AcCoA node. The overall reaction for hybrid EMP was obtained (**eq. 4**). (**B**) Stoichiometric comparison of hybrid EMP with either native EMP or NOG for pyruvate or acetyl-CoA biosynthesis in terms of ATP, inorganic phosphate, NADH, and CO_2_. Succinyl-CoA; SuccCoA.

To obtain the overall reaction for the hybrid EMP pathway, we made the assumption (**eq. 1–1)** that 2 G3P molecules, which is an intermediate of the NOG pathway, obtained from 2 glucose molecules are hijacked by glyceraldehyde 3-phosphate dehydrogenase and used during the pay-off phase of the EMP pathway due to the xylulose 5-phosphate activity and as a result, 3 AcP molecules can be produced from 2 glucose (**eq. 1–1)** to (**eq. 1–2)**. Then, the AcP-AcCoA node can be used for the overall reaction (**eq. 3**). We found that there were two possible routes, either via phosphate acetyltransferase or acetate kinase/acetyl-CoA:acetate CoA transferase. Thus, we derived the overall reaction for the hybrid EMP pathway, depending on the carbon flux ratio at the AcP-AcCoA node (**eq. 4**):

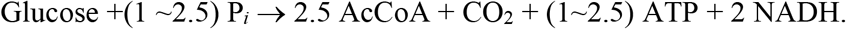

Although there is no production of CO_2_ per glucose via NOG during acetyl-CoA biosynthesis, hybrid EMP generates 1 CO_2_ per glucose while EMP produces 2 CO_2_ per glucose (**Fig. 2B**). It is possible to preserve half the CO_2_ produced using hybrid EMP compared to native EMP. In case of pyruvate synthesis from glucose, 1.5 CO_2_ per glucose can be produced using NOG and a single ATP is required^10^. Whereas in the hybrid EMP, ATP/NADH can be generated without CO_2_ loss during pyruvate synthesis and 1.5 mole acetate can be obtained instead of CO_2_.

Since Pkt plays an important role in NOG, the P_*i*_ levels must be taken into consideration along with ATP for the overall reaction. Although higher P_*i*_ supply is required to run the hybrid EMP than either native EMP or NOG, the net ATP produced is also higher than NOG. Unlike NOG, hybrid EMP is still able to produce NADH as reducing power while CO_2_ production is reduced. Compared to EMP, reduction of 1 CO_2_ resulted in a reduction of 2 NADH in hybrid EMP. Thus, we aimed to elucidate the hybrid EMP-mediated metabolism that can support aerobic cell growth with lesser CO_2_ production and higher acetyl-CoA levels with respect to P_*i*_ over the native EMP. We designed the reconstruction of the hybrid EMP with NOG in an industrial L-lysine producer, *Corynebacterium glutamicum* as a sample organism, to reduce the CO_2_ production without compromising the cell growth in minimal glucose medium.

### Blocking the preparatory phase of EMP and combining with NOG pathway

To construct the *C. glutamicum* strains with the hybrid EMP pathway (**Fig. 3A**), we first deleted *pfkA* to block the biosynthesis of fructose bisphosphate from fructose 6-phosphate (strain YL1; Supplementary Table S1, Fig. S1) to block the preparatory phase of EMP. Compared to WT pZ8-1 (an empty vector) as a control, YL1 pZ8-1 exhibited strong growth inhibition in minimal glucose medium (4.2 ± 0.26 of OD_600_ in 72 h) (**Fig. 3B**). Then, the second phase of the hybrid EMP pathway can be initiated in the YL1 strain by introducing the NOG pathway. Heterologous Pkt from *Bifidobacterium adolescentis*^9,12^, which can hydrolyze F6P (Fpk) as well as xylulose 5-phosphate (Xpk), was expressed in YL1 (strain YL1 pZ8-fxpk). However, there was no improvement in the growth rate compared to YL1 pZ8-1. Although glucose can be transported via the PTS system and glucose 6-phosphate can be metabolized in the EMP pathway, there can be an imbalance in the net ATP in the hybrid EMP system. For efficient ATP utilization by the YL1 pZ8-fxpk strain, an additional enzyme, a glucokinase (Glk; native *glk*)), was co-expressed and as a result, the growth rate of YL1 pNOG1 was completely recovered at similar OD_600_ compared to WT pZ8-1. However, the sole expression of *glk* in YL1 pZ8-glk could not enhance the growth rate. Therefore, we concluded that the synthetic activities of both Pkt and Glk are required for functional hybrid EMP pathway in YL1 pNOG1 in minimal glucose medium. To accelerate the growth of the strain with hybrid EMP, native transaldolase (Tal) was also overexpressed in YL1 pNOG2 as Tal activity limits the pentose catabolism in *C. glutamicum* rather than transketolase (Tkt)^12^. Therefore, YL1 pNOG2 was used as the reference strain for the hybrid EMP pathway in *C. glutamicum*.

**Figure 3.**
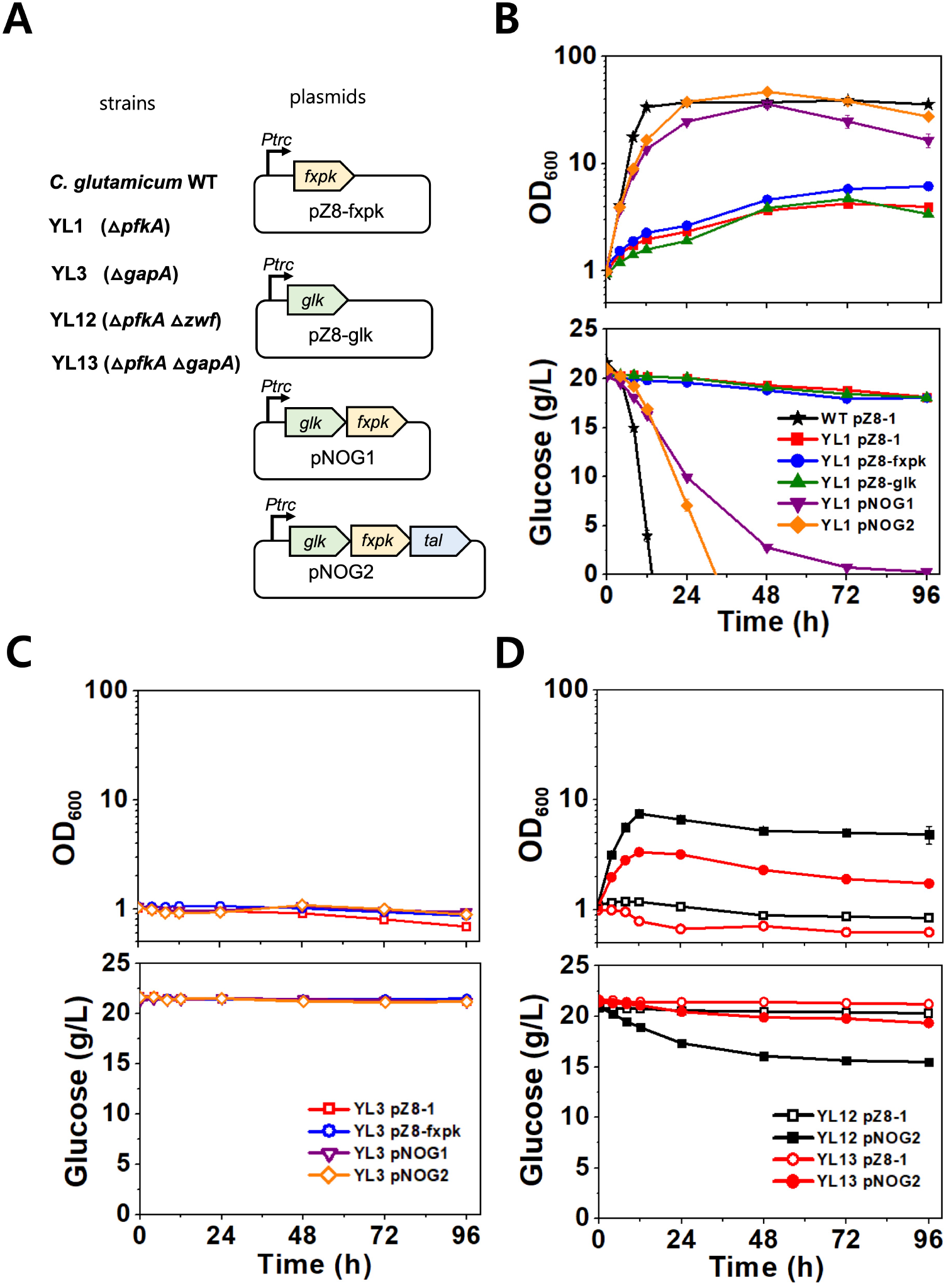
Growth and glucose consumption in engineered *C. glutamicum* strains on hybrid EMP pathway. (**A**) Names of strains and plasmids used for the hybrid EMP pathway were described. The details were shown in Supplementary Table S1. (**B** – **D**) Growth (optical density at 600 nm, upper panel) and glucose consumption (g/L, lower panel). The strains were cultured in CgXII minimal medium (50 mL in 250 mL baffled Erlenmeyer flasks) with 2% (w/v) glucose as the sole carbon source. Data represent mean values of at least triplicated cultivations, and error bars represent the standard deviations.

Single deletion of *gapA* encoding for glyceraldehyde 3-phosphate dehydrogenase in *C. glutamicum* completely abolishes cell growth in minimal glucose medium (**Fig. 3C**, Supplementary **Fig. S1**). None of the plasmids used for establishing hybrid EMP in YL3 supported sufficient cell growth in minimal glucose medium except in YL13 pNOG2 (3.35 ± 0.13 of OD_600_ in 12 h) (**Fig. 3D**).

Although the corynebacterial K_m_ value has not identified, the *E. coli* K_m_ value of PfkA is a 1000-fold lower than that of Fxpk (Pkt from *Bifidobacterium*) when F6P is used as the competing substrate^13,14^. Thus, F6P could not be converted by Fxpk due to its lower activity in the presence of PfkA (YL3 pNOG2), resulting in no cell growth. However, YL13 pNOG2 lacking PfkA could convert F6P to AcP and erythrose-4-phosphate (E4P), resulting in significant cell growth. This result implicated the synthetic NOG pathway as the first phase of the hybrid EMP was indeed active in *C. glutamicum*.

Unlike the *E. coli* NOG strains^10^, adaptive laboratory evolution of *C. glutamicum* does not require growth in minimal glucose medium. To investigate whether NOG is an active pathway in the hybrid EMP pathway, *zwf* (encoding for glucose 6-phosphate dehydrogenase) was deleted to block the oxidative phosphate pathway that was only a possible catabolic pathway in YL1. As a result, YL12 pZ8-1 did not grow on glucose in minimal medium. However, YL12 pNOG2 exhibited considerable cell growth in minimal glucose medium (7.48 ± 0.07 of OD_600_ in 12 h), although complete glucose consumption was not achieved (**Fig. 3D**). Therefore, we conclude that replacing PfkA with Glk and utilizing active NOG in the hybrid EMP provides an alternative glycolytic pathway.

### Identifying the metabolic fate of acetyl phosphate in the hybrid EMP strains metabolizing glucose

Since NOG pathway provides balanced carbon rearrangement from F6P to AcP in the hybrid EMP, we investigated how AcP can be further metabolized to acetyl-CoA in YL1 pNOG2 with hybrid EMP. There are two possible routes at the AcP-AcCoA node that exist in bacteria: route (i) direct conversion to acetyl-CoA by Pta (encoded by *pta*), and route (ii) two-step conversion: (**a**) conversion of AcP to acetate and ATP by acetyl kinase (encoded by *ackA*) and (**b**) conversion of acetate and CoA to acetyl-CoA and ATP by either succinyl-CoA:acetate CoA-transferase (encoded by *actA*) or acetyl-CoA synthetase (encoded by *acs*). As there is lack of Acs activity in *C. glutamicum*, only activities of AckA and ActA were taken into consideration in the second route (**Fig. 2B**).

First, we deleted each gene at the AcP-AcCoA node (*pta*, *actA*, and *ackA*) in the strain YL1, and the resultant strains were named YL14, YL15, and YL16, respectively. All mutants harboring pNOG2 exhibited slower growth rate and glucose consumption rate, compared to YL1 pNOG2 (**Fig. 4**). YL14 pNOG2 (Δ*pta*; Δroute i) and YL16 pNOG2 (Δ*ack*; Δroute ii-**a**) showed stronger growth inhibition and slower glucose consumption than either YL12 pNOG2 or YL15 pNOG2 (Δ*actA*; Δroute ii-**b**). Thus, we concluded that each route is active but not redundant in hybrid EMP. In case of YL15 pNOG2, the cell growth was better than the other strains. This could be due the fact that the production of ATP and P_*i*_ was not compromised in YL15 pNOG2 due to the AckA activity. However, YL16 pNOG2 and YL14 pNOG2 showed lower ATP levels than YL12 pNOG2 (**Fig. 4A**), as the consequence of the imbalance of ATP and P_*i*_. Thus, we concluded that YL1 pNOG2 utilized both flexible routes (i) and (ii) at the AcP-AcCoA node in the hybrid EMP although both routes are energy-expense reactions. The hybrid EMP pathway balanced both ATP and P_*i*_ usages from the routes against glucokinase and phosphoketolase reactions.

**Figure 4.**
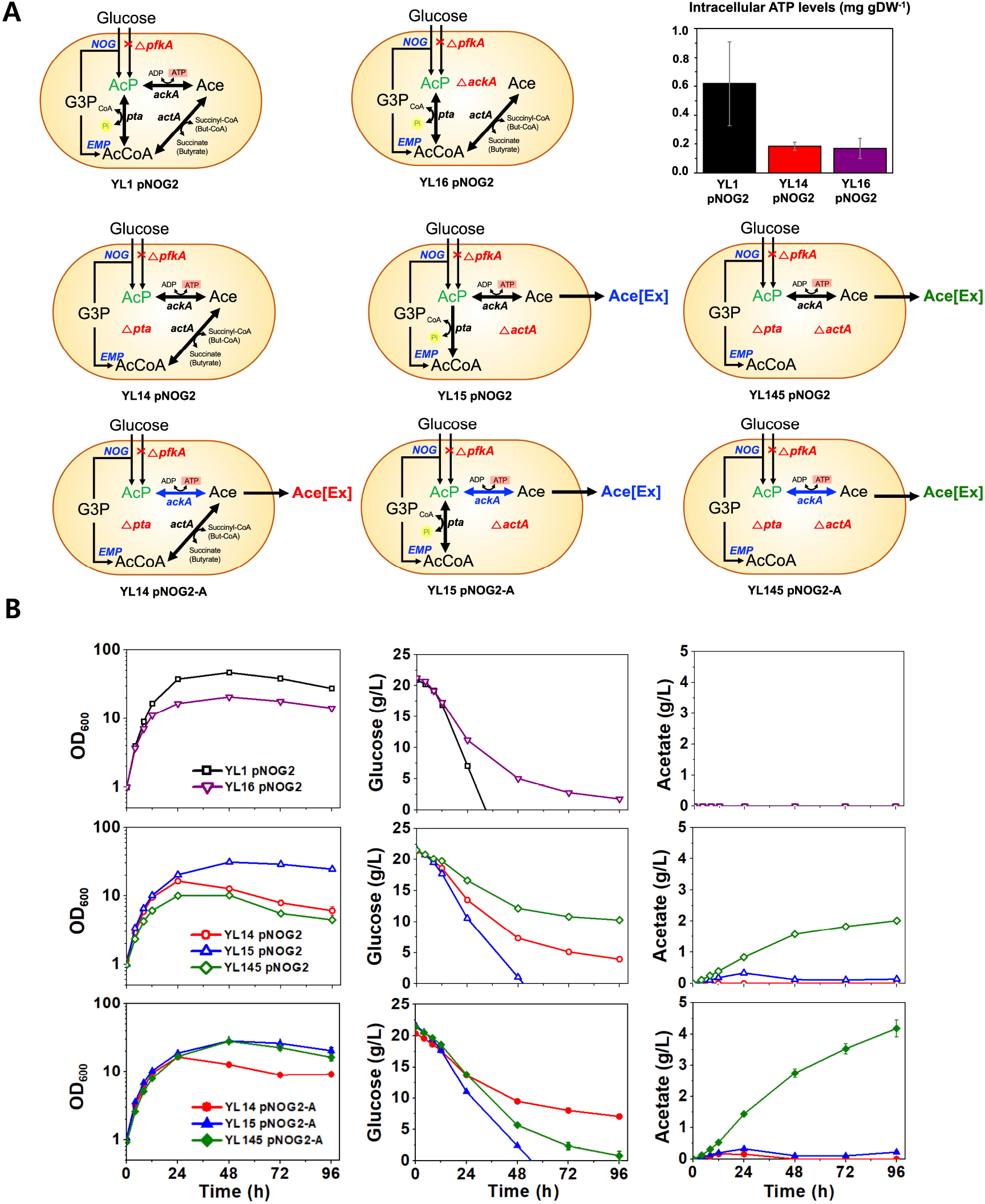
Metabolic fate of acetyl phosphate generated from hybrid EMP in the *C. glutamicum*. (**A**) Scheme of the metabolic pathways for the hybrid EMP strains were described with the genotypes. Derivative strains from a hybrid EMP strain (YL1 pNOG2) were investigated to identify the fate of acetyl phosphate (AcP) generated from the NOG pathway in *C. glutamicum*. Strains were constructed by using Coryne-CR12-Del. Blue arrows represent overexpressed reactions. Intracellular ATP levels were measured using the bioluminescence assay. Data represent mean values of at least triplicated cultivations, and error bars represent the standard deviations. (**B**) Measurement of the growth, glucose, and acetate from the strains. Color and symbols represent strains used in this study: YL1 pNOG2 (black; open square), YL14 pNOG2 (red; open circle), YL14 pNOG2-A (red; solid circle), YL15 pNOG2 (blue; open triangle), YL15 pNOG2-A (blue; solid triangle), YL145 pNOG2 (green; open diamond), YL145 pNOG2-A (green; solid diamond), and YL16 pNOG2 (purple; open inverted triangle). See the Table S1 for the details. Data represent mean values of at least triplicated cultivations, and error bars represent the standard deviations.

Interestingly, we found that during aerobic cultivation of YL15 pNOG2, a small amount of acetate was secreted into the medium and glucose was depleted within 48 h. Thus, we assumed that acetate accumulates in the cell and is slowly converted to AcP, which is then converted to acetyl-CoA. However, 2.0 ± 0.01 g/L acetate was secreted when we deleted *pta* in YL15 pNOG2 (YL145 pNOG2). The reason behind this could be the increased AcP level in YL145 pNOG2, which forced it to secrete more acetate than YL15 pNOG2. Imbalance due to limited ATP and P_*i*_ generation via Ack and Glk resulted in incomplete glucose consumption. When we overexpressed *ackA* in the AcP-AcCoA mutants, only YL145 pNOG2-A exhibited growth improvement and glucose consumption by secreting increased amounts of acetate (4.18 ± 0.27 g/L). Hence, excessive ATP generation provide by Ack supplied the limited P_*i*_ required for YL145 pNOG2. In parallel, Pta was overexpressed in YL16 pNOG2-P lacking Ack, and cell growth was slightly increased to due to increased conversion of AcP to acetyl-CoA and P_*i*_ (**Fig. S2**). Therefore, we proposed that while the route (i) was a major path at AcP-AcCoA node, route (ii-**a, b)** was also essential to support complete aerobic cell growth in *C. glutamicum* with hybrid EMP.

### Link between sugar and phosphate metabolism via trehalose biosynthesis

We noticed that the cell growth was lower in the hybrid EMP strains, including YL1 pNOG2, than in WT pZ8-1 when glucose was completely depleted and none of the organic acids were detected in the hybrid EMP strains. However, we observed an unexpected increase in the disaccharide peaks over time when we analyzed the medium samples for the engineered strains. No disaccharides were detected for the WT strains in the minimal glucose medium. To identify the unknown sugars, we performed enzymatic assays using authentic standards and diluted supernatant samples (**Fig. S3**), which ultimately revealed the presence of trehalose synthesis and secretion. Then, we analyzed the all the samples and calculated the concentrations as well as the specific production of trehalose (**Fig. 5B**) and observed higher specific trehalose production in all the mutants of the hybrid EMP strain compared to the parental strain (YL1 pNOG2) except in the YL15 pNOG2 strain. YL15 pNOG2 has both native Pta and Ack activities that results in the production of 1 P_*i*_ or 1 ATP from 1 AcP. YL14 pNOG2, which lacks native Pta, exhibited the highest specific trehalose production among the engineered strains. This could be due to the fact that ATP generation from accumulated AcP is beneficial for cellular fitness. Since hybrid EMP needed both routes (i) and (ii) at the AcP-AcCoA node for cell growth, imbalances resulting from biased routes may affect the cell growth. Thus, we reasonably assumed that there might be a link between the trehalose biosynthesis pathway and AcP catabolism.

**Figure 5.**
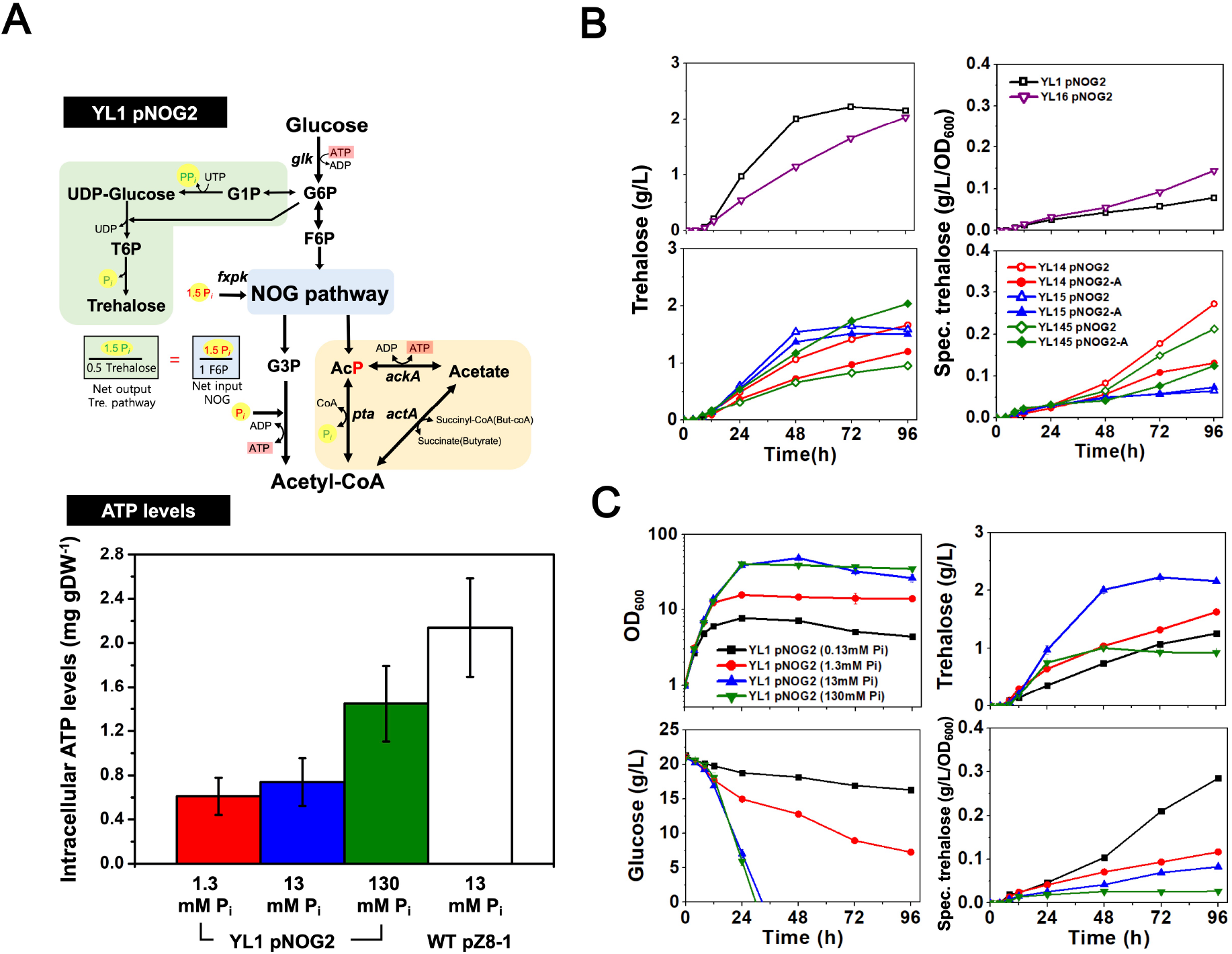
Replenishing inorganic phosphate sources by a trehalose synthesis on the hybrid EMP. (**A**) a metabolic pathway of trehalose biosynthesis and hybrid EMP pathway in YL1 pNOG2. The green highlighted path indicates trehalose biosynthesis pathway via UDP-glucose, generating a net 1.5 mole inorganic phosphate per 1 mole trehalose produced from 2 mole of glucose 6-phosphate. The blue highlighted path indicates the NOG pathway, requiring a net 1.5 mole inorganic phosphate per 1 mole of fructose-phosphate as substrate. AcP as an output from NOG can be converted to acetyl-CoA via flexible AcP-acetate node. For the intracellular ATP measurement, YL1 pNOG2 was cultivated in CgXII medium with 2% (w/v) glucose with various inorganic phosphate sources (1.3 mM, red; 13 mM, blue as the normal CgXII, 130 mM, green) and WT pZ8-1 was cultivated with 13 mM P_i_ as control (white bar). Intracellular ATP levels were measured using the bioluminescence assay. The hybrid EMP strains were analyzed for trehalose concentration and specific trehalose production were calculated. Supernatants from the hybrid EMP strains were analyzed for quantification of trehalose using HPLC after the enzymatic confirmation (**Fig. S3**). Data represent mean values of at least triplicated cultivations, and error bars represent the standard deviations. (**B**) Measurement of trehalose in the medium and specific trehalose (g/L/OD_600_) was calculated. Color and symbols represent strains used in this study: YL1 pNOG2 (black; open square), YL14 pNOG2 (red; open circle), YL14 pNOG2-A (red; solid circle), YL15 pNOG2 (blue; open triangle), YL15 pNOG2-A (blue; solid triangle), YL145 pNOG2 (green; open diamond), YL145 pNOG2-A (green; solid diamond), and YL16 pNOG2 (purple; open inverted triangle). Data represent mean values of at least triplicated cultivations, and error bars represent the standard deviations. (**C**) Growth (optical density at OD_600_), glucose consumption (g/L), trehalose secretion (g/L), specific trehalose production (g/L/OD_600_). YL1 pNOG2 was cultured in CgXII medium (50 mL in 250 mL baffled Erlenmeyer flasks) with 2% (w/v) glucose as the sole carbon source and various inorganic phosphate sources (13 mM as control, blue; 0.13 mM, black; 1.3 mM, red; 130 mM, green). Data represent mean values of at least triplicated cultivations, and error bars represent the standard deviations.

When we calculated the metabolic stoichiometry to understand how the hybrid EMP strains benefit from production of 1 trehalose from 2 glucose 6-phosphate (or fructose 6-phosphate), we observed that 3 P_*i*_ can be released by synthesis of 1 trehalose molecule via UDP-glucose^15^. As 1.5 P_*i*_ is necessary for NOG of a F6P molecule (**Fig. 2A**), half of trehalose synthesis can supply this to the NOG in hybrid EMP. Based on the fact that hybrid EMP requires higher P_*i*_ than native EMP, we tested with various P_*i*_ concentrations to evaluate the effects on trehalose biosynthesis in hybrid EMP and as a result, we discovered a link between sugar and phosphate metabolism. As a result, the specific production of trehalose was decreased as more P_*i*_ was utilized for the hybrid EMP (**Fig. 5C**). This conclusively proves that trehalose biosynthesis not only plays a role as an osmoprotectant and stress protectant^16,17^ but also as an internal modular that acts in response to limited internal P_*i*_ pools. Thus, we found that the metabolic link between carbon metabolism and phosphorus metabolism is indeed tightly interconnected in a balanced metabolic state^7^. Evaluation of the hybrid EMP also showed that carbon and phosphorus metabolism are interconnected during high P_*i*_ requirement.

### Fermentation Profile of the Hybrid EMP Strains

Based on the overall equations (native EMP and hybrid EMP), we proposed that 50% less CO_2_ and 25% more acetyl-CoA will be produced per glucose in the hybrid EMP (**Fig. 2B**). To demonstrate the capability of the hybrid EMP to reduce CO_2_ production and increase the acetyl-CoA levels, a controlled bioreactor was used to continuously monitor the CO_2_ production with dissolved oxygen levels for YL1 pNOG2 with WT pZ8-1 as the control. In addition, cell growth, glucose consumption, and specific intracellular acetyl-CoA levels were measured from the collected samples. As a result, we found that the final optical density (48 ± 1.2 at OD_600_) of YL1 pNOG2 was approximately the same as the optical density (50.1 ± 0.3 at OD_600_) of WT pZ8-1, although the growth rate of YL1 pNOG2 (μ, 0.22 ± 0.01 h^−1^) was half of the WT pZ8-1 (μ, 0.41 ± 0.004 h^−1^) growth rate (**Fig. 6A**). Further, we obtained 2.2 ± 0.01 g/L of trehalose from the YL1 pNOG2 culture, as shown in the flask culture. As expected, the total CO_2_ production from the YL1 pNOG2 culture was 82.6% of that of the WT pZ8-1 culture until glucose was completely depleted. As the net CO_2_ evolved during hybrid EMP should be more than that during the pyruvate dehydrogenase reaction, the fact that we observed approximately 20% reduction in CO_2_ loss in the hybrid EMP compared to the native EMP was impressive. Although the glucose uptake system was not manipulated in YL1 pNOG2, the specific glucose uptake rate (0.89 ± 0.001 mmol/gDW/h) of YL1 pNOG2 was lower than that of the WT pZ8-1 (2.55 ± 0.01 mmol/gDW/h). This could be due to the redistribution of sugar metabolism in the hybrid EMP due to the activity of Glk, which might have weakened the thermodynamic driving forces from glucose under P_*i*_ limited conditions^18^. Subsequently, specific acetyl-CoA levels were measured, and a 19% increase was observed in YL1 pNOG2 over WT pZ8-1 (**Fig. 6B**). Thus, we demonstrated that glucose metabolism can occur with less CO_2_ and more acetyl-CoA production in engineered strains via hybrid EMP compared to native EMP.

**Figure 6.**
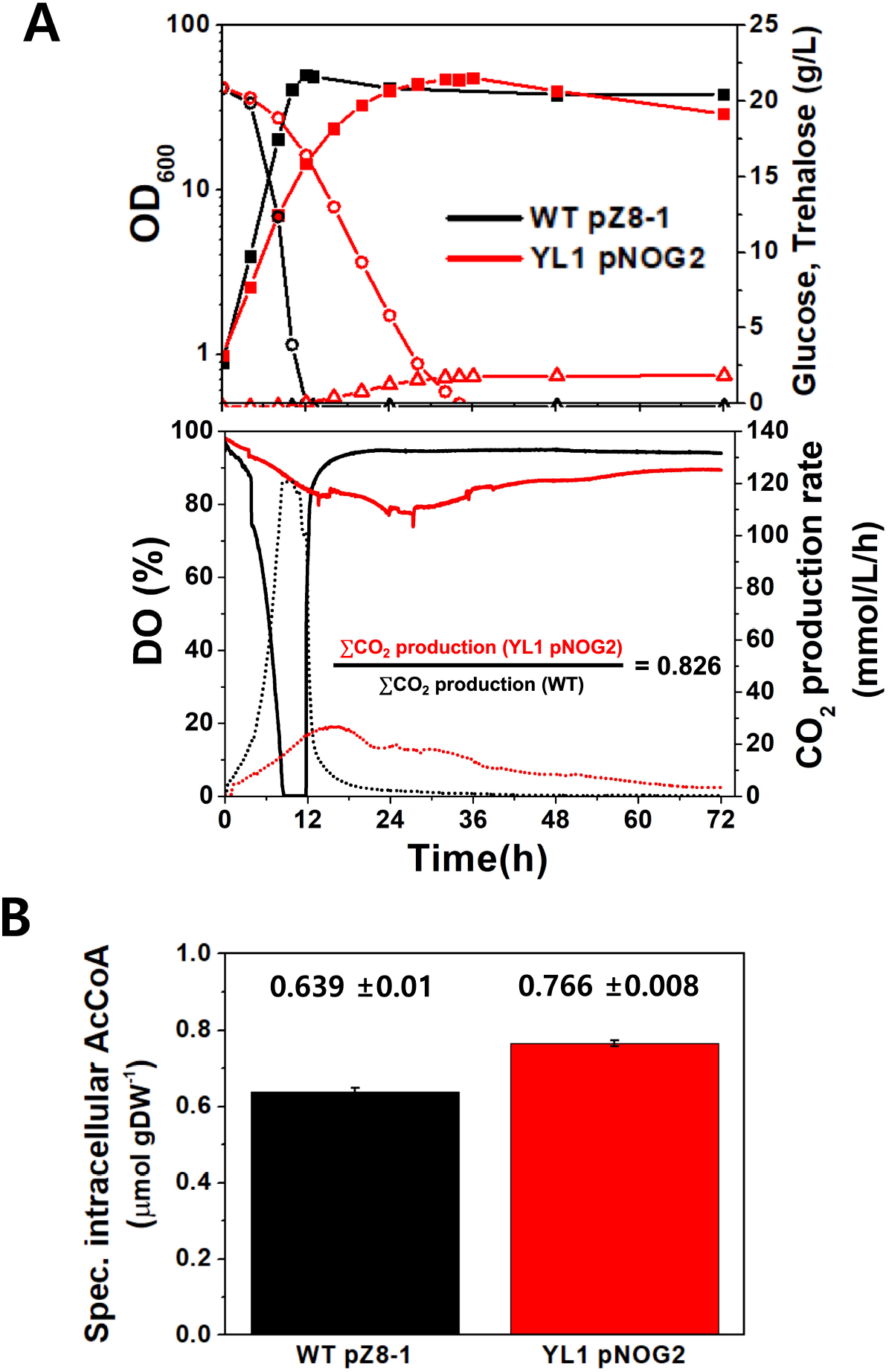
A decrease of CO_2_ evolution and an increase of acetyl-CoA on the hybrid EMP. (**A**) Measurement of the growth (solid square), glucose (open circle), trehalose (open triangle) dissolved oxygen (DO, %), and CO_2_ production rate (mmol/L/h) from either WT pZ8-1 (black; a control) or YL1 pNOG2 (red). Relative CO_2_ production was calculated with respect to the strains. (**B**) Specific intracellular acetyl-CoA (AcCoA) were measured using the Acetyl CoA Fluorometric Assay Kit. Data represent mean values of at least triplicated cultivations, and error bars represent the standard deviations. An OD_600_ of 1 is converted to 0.3 g cell dry weight per liter.

## Discussion

In this study, we developed a hybrid EMP pathway by rewiring it with a synthetic NOG pathway and modifying the native EMP pathway. We constructed an engineered *C. glutamicum* strain with the hybrid EMP. After confirming that the engineered *C. glutamicum* with hybrid EMP has approximately similar amount of biomass as the *C. glutamicum* WT in the minimal glucose medium, the AcP–acetyl-CoA nodes were deciphered to understand the complete carbon metabolism to generate acetyl-CoA. As a result, a metabolic link between the carbon and phosphorus metabolism was revealed in the hybrid EMP.

Starvation of cells for inorganic phosphate is detrimental to cell growth ^7^. Unlike the native EMP pathway, the hybrid EMP pathway requires additional 0.5 P_*i*_ per glucose. Thus, the cells must regulate their internal phosphate sources for cell growth until additional P_*i*_ is supplemented. We revealed that there is a metabolic link between the AcP–acetyl-CoA node and trehalose biosynthesis and this was responsible for regulating internal phosphate sources. Similarly, a previous study has shown that external phosphate starvation was linked to glycogen metabolism at the metabolite level, resulting in high glycogen levels in the P_*i*_ limited medium^19^. In diatoms, which are unicellular photosynthetic phytoplankton, phosphate limitation triggers accumulation of extracellular polysaccharides with UDP-sugars as the substrates^8^. In addition, sugar-mediated regulation via ADP-glucose pyrophosphorylase has been uncovered as the P_*i*_ sequestering mechanism in tobacco seedlings^20^. Moreover, trehalose biosynthesis was revealed to be a futile cycle for restoring the balanced state of glucose when inorganic phosphate sources and ATP levels are provided^7^. Inorganic phosphate can be sensed by specific phosphate carriers, which can act cooperatively with sugar metabolism by activating protein kinases in *Saccharomyces cerevisiae*^21^. Rapid phosphate signaling was found to be dependent on the sensing of glucose by a glucose-phosphorylation dependent system. Thus, activation of trehalose 6-phosphate synthase (*otsA*) and trehalose phosphatase (*otsB*) in the hybrid EMP pathway might be involved in the internal phosphate signaling and phosphate uptake via phosphate transporters or a membrane H^+^-ATPase. Genetic regulation for both phosphate signaling and the metabolic link has not been identified yet in *C. glutamicum*. Specific trehalose exporters responsible for secretion remain unclear in *C. glutamicum*^22^. None the less, carbon metabolism can be flexible at the metabolite or protein level in response to internal or external environmental changes of phosphate sources via trehalose biosynthesis.

*Bifidobacterium*, an anaerobic gram-positive bacilli, found in nature have a bifid shunt that converts hexose to acetic acid and lactic acid and produces a high net of 2.5 ATP per glucose, which is more effective than the EMP^9,23,24^. The bifid shunt has also evolved in heterofermentative lactic acid bacteria to adapt to various environments by secreting lactate and acetate. Although the hybrid EMP constructed in this study also incorporates the bifid shunt activities of an oxidative EMP and Pkt, as the engineered *C. glutamicum* with the hybrid EMP possesses the key Glk activity, it cannot secrete both lactic acid and acetic acid and hence under aerobic or microanaerobic condition, has the advantage of highly efficient energy generation with a maximum of 2.5 ATP produced per glucose molecule. In addition, the net ATPs produced per glucose in engineered bacteria can be controlled by controlling the initial inorganic phosphate concentration and regulating the AcP–acetyl-CoA node.

The hybrid EMP pathway produces a net of 2 NADH per glucose, compared to no NADH produced by the synthetic NOG pathway in order to produce acetyl-CoA (**Fig. 2B**). Thus, an oxidative pentose phosphate pathway and oxidative conversion of glyceraldehyde-3-phosphate in the pay-off pathway are possible from glucose as the sole carbon and energy source in the engineered strains with hybrid EMP. This characteristic catabolic mechanism is favorable for the synthetic NOG that requires external reducing equivalents such as H2 or formic acid to carry out sugar metabolism for CO_2_ fixation^10^.

In conclusion, the hybrid EMP pathway allows development of microbial strains capable of reduced CO_2_ and enhanced acetyl-CoA production in the balanced phosphate state without losing any NADH. By controlling the initial inorganic phosphate concentration on hybrid EMP, the levels of acetyl-CoA–derived products can be enhanced with higher carbon yields, which will provide an alternative mode for microbial physiology as well as an opportunity for potential industrial applications.

## Materials and Methods

### Chemicals and Reagents

All of the chemicals used in this study was purchased at Sigma-Aldrich (St. Louis, MO) unless otherwise mentioned. High fidelity polymerase (PrimeSTAR GXL, Takara Bio Inc, Shiga, Japan) was used for PCR. All the PCR products and plasmids were purified and extracted using the Dyne Plasmid Miniprep Kit (DYNEBIO, Gyeonggi, Korea) and Expin Combo GP (GeneAll Biotechnology, Seoul, Korea). All the oligonucleotides were purchased from GenoTech (Daejeon, Korea) and DNA sequencing was performed by Macrogen (Seoul, Korea).

### Strains and growth conditions

*E. coli* DH5α^25^ and *C. glutamicum* ATCC 13032 (WT) were used in this study (Supplementary Table S1). *E. coli* strains were grown in Lysogeny Broth (LB) medium (10 g/L tryptone, 5 g/L yeast extract, and 5 g/L NaCl) at 37°С on a rotary shaker at 200 rpm. When required, the medium was supplemented with 50 μg/mL of kanamycin. For the construction of the hybrid EMP pathway in *C. glutamicum*, the target chromosomal genes were inactivated by using Coryne-CR12-Del (<u>Coryne</u>bacterial <u>C</u>RISPR-based <u>R</u>ecombineering using Cas<u>12</u>a to <u>del</u>ete 100 bp in the coding sequence)^26,27^. The plasmids for gene amplification were introduced into *C. glutamicum* by electroporation, and strain validation was performed using colony PCR and DNA sequencing^28^. *C. glutamicum* ATCC 13032 and its derivatives strains were pre-cultivated in brain heart infusion-supplemented (BHIS) medium overnight and then incubated aerobically in CgXII defined medium (50 mL media in a 250 mL baffled Erlenmeyer flask) containing 2% (w/v) glucose as the sole carbon source at 30°C on a rotary shaker at 120 rpm^12,29^. When required, the medium was supplemented with 25 μg/mL of kanamycin and/or spectinomycin.

### Plasmid construction

DNA manipulation and cloning for pathway construction were performed according to standard protocols^30^. The primers used in this study are listed in *SI Appendix* Table S2. crRNA plasmids carrying crRNAs were constructed from pJYS2_crtYf (Supplementary Table S3). Target crRNAs for recombineering were cloned into the standard vector using a primer containing a target-specific protospacer region. A 59 bp single-stranded oligodeoxynucleotide (lagging strand ssODN) was designed to delete 100 bp of the coding sequence region without causing any translational initiation or frameshift mutation (Supplementary Table S2). The crRNA and ssODN were co-transferred into competent Corynebacterial cells harboring pJYS1Ptac. To overexpress the genes, the *fxpk* gene from *Bifidobacterium adolescentis*^9,12^ was synthesized (Genscript, USA) with codon-optimization for *C. glutamicum*. The native genes (*glk*, *tal*, *ackA*, and *pta*) were amplified from the genomic DNA of *C. glutamicum* and cloned into the pZ8-1 vector (Supplementary Table S1).

### Measurement of metabolite levels

Glucose, trehalose, and organic acids in the supernatant were quantified using high-performance liquid chromatography (HPLC) as described previously^31^. Briefly, the culture supernatant was passed through a syringe filter (pore size = 0.45 μm). The concentrations of glucose and organic acids were detected by a HPLC (Agilent 1260, Waldbronn, Germany) equipped with a refractive index detector (RID), and a Hi-Plex H, 7.7 × 300 mm column (Agilent Technologies) under the following conditions: sample volume of 20 μL, mobile phase of 5 mM H_2_SO_4_, a flow rate of 0.6 mL/min, and column temperature of 65°C. To identify the potential secreted disaccharides (trehalose or maltose), enzymatic assay was applied to the diluted supernatant using either trehalase (1 U/mg; Sigma No. T8778) or α-glucosidase (Bottle 5 in the Maltose assay kit; Megazyme Inc., Wicklow, Ireland) (**Fig. S3**). For the ATP assay, the intracellular ATP were measured using the enzymatic assay kit (ATP Bioluminescence Asssay Kit HS II, Sigma No. 11699709001, Roche, Switzerland) after the cell disruption of the corynebacterial strains. Briefly, the culture condition for the intracellular ATP measurement was identical to the conditions for other metabolites. The cells were harvested by centrifugation at 5,000 x *g* for 5 min after the optical densities of the cells reached at the exponential phase. The cell pellets were disrupted by glass bead-beating. Then, the cell extracts were used for the bioluminescenct ATP assay and all the procedures were performed based on the manufacturer’s instructions.

### Measurement of enzymatic acetyl-CoA

Recombinant strains were cultivated in CgXII medium containing 2% (w/v) glucose. Briefly, a 1 mL-culture sample from the exponential growth phase was centrifuged (3381 × *g*), and the cells were washed with a Tris-based buffer (pH 6.3). The cell pellet was normalized to the optical density of 1 and was re-suspended in 1 mL of the assay buffer (Acetyl CoA Fluorometric Assay Kit; BioVision, Milpitas, CA) along with 0.1 mm diameter glass beads. After a glass-bead beating, the supernatant (500 μL) was collected and used for the Acetyl CoA Fluorometric Assay Kit (BioVision, Milpitas, CA). The fluorescence was measured using Infinite® M Nano microplate reader (Ex/Em:535/587; Tecan AG, Switzerland). All the procedures were performed based on the manufacturer’s instructions.

### Fermentation cultivation

Batch fermentation was conducted in a 5 L jar fermenter (MARADO-05S-XS; CNS Inc., Daejeon, South Korea) containing 2 L culture medium. The seed culture was prepared in a 250 mL Erlenmeyer flask containing 100 mL BHIS medium at 30°С on a rotary shaker at 120 rpm. The seed culture was transferred to a fermenter at an OD_600_ of 1. The fermenter contained 2 L CgXII medium containing either 13 mM or 130 mM inorganic phosphate (P_i_) and 2% (w/v) glucose as the sole carbon source. P_i_ was added in the form of KH_2_PO_4_ and K_2_HPO_4_. The main culture was maintained under 25 μg/mL of kanamycin. The pH of the culture was adjusted to 7 by adding either 5 N NaOH or 6 N HCl. Filtered air was constantly supplied at 1 L/min (0.5 [vvm]), and the dissolved oxygen (DO) level was monitored at constant 400 rpm. CO_2_ production rate (mmol/L/h) was calculated by measuring the CO_2_ percentage (%) by digitally recording every per min using a CO_2_ analyzer (Q-S153, Qubit systems, Kingston, ON, Canada). Then, the total CO_2_ production rates (ΣCO_2_) were calculated by integrating the area of the curve until the initial glucose was completely depleted.

## Supporting information

Supplmentary Tables and Figues

## Additional information

**Supplementary Information** accompanies this paper at

## ACKNOWLEDGMENTS

This work was supported by the Technology Innovation Program (No. 20000158, 20000679), funded by the Ministry of Trade, Industry and Energy, and the Individual Research Program (2020R1F1A1048292) through the National Research Foundation of Korea, Republic of Korea.

## Author contributions

Y.L., H.J.C., and H.M.W. designed research; Y.L. and H.J.C. performed research; Y.L., H.J.C., and H.M.W. analyzed data and wrote the paper.

