## Supplementary material for "A hybrid Embden–Meyerhof–Parnas pathway provides a synthetic link between sugar and phosphate metabolism": Supplmentary Tables and Figues

**Supplementary Table S1.** Bacterial strains and plasmids used in this study.

| Strain or plasmid | Relevant characteristics | References |
|---|---|---|
| Strains | | |
| *E. coli* DH5α | F−(80d ***lac***Z M15) (*lac*ZYA-argF) U169 *hsd*R17(r− m+) *rec*A1 *end*A1 *rel*A1 *deo*R96 | (1) |
| *C. glutamicum* WT | ATCC 13032, wild-type strain | ATCC |
| YL1 | *C. glutamicum* ATCC 13032, Δ*pfkA* (CDS 100 bp deletion) | This study |
| YL2 | *C. glutamicum* ATCC 13032, Δ*zwf* (CDS 100 bp deletion) | Lab stock |
| YL3 | *C. glutamicum* ATCC 13032, Δ*gapA* (CDS 100 bp deletion) | This study |
| YL12 | YL1, Δ*zwf* (CDS 100 bp deletion) | This study |
| YL13 | YL1, Δ*gapA* (CDS 100 bp deletion) | This study |
| YL14 | YL1, Δ*pta* (CDS 100 bp deletion) | This study |
| YL15 | YL1, Δ*actA* (CDS 100 bp deletion) | This study |
| YL16 | YL1, Δ*ackA* (CDS 100 bp deletion) | This study |
| YL145 | YL14, Δ*actA* (CDS 100 bp deletion) | This study |
| Plasmids | | |
| pZ8-1 | pHM1519, $Km^r$, $P_{tac}$, *E. coli*-*C. glutamicum* shuttle vector | (2) |
| pZ8-fxpk | pZ8-1 vector carrying the *fxpk* gene of *B. adolescentis* codon-optimized for *C. glutamicum* | This study |
| pZ8-glk | pZ8-1 vector carrying the *glk* gene(cg2399) of *C. glutamicum* | This study |
| pZ8-tal | pZ8-1 vector carrying the *tal* gene(cg1776) of *C. glutamicum* | Lab stock |
| pNOG1 | pZ8-1 vector carrying the *glk* gene(cg2399) of *C. glutamicum* and *fxpk* gene of *B. adolescentis* codon-optimized for *C. glutamicum* | This study |
| pNOG2 | pNOG1 vector carrying the *tal* gene(cg1776) of *C. glutamicum* | This study |
| pNOG2-A | pNOG2 vector carrying the *ackA* gene(cg3047) of *C. glutamicum* | This study |
| pJYS1Ptac | $pBL1^{ts}$ *ori*$V_{C.g.}$ $Km^r$ pSC101 *ori*$V_{E.c.}$ PlacM-*Fn*Cpf1 $P_{tac}$-RecT *lacI*$^q$ | (3) |
| pJYS2_crtYf | *rep ori*$V_{C.g.}$ $Sp^r$ pMB1 *ori*$V_{E.c.}$ $P_{j23119}$-crRNA-crtYf targeting the *crtYf* gene | (3) |
| pcrRNA-pfkA | pJYS2 derivative, $P_{j23119}$-crRNA-pfkA targeting the *pfkA* gene | This study |
| pcrRNA-gapA | pJYS2 derivative, $P_{j23119}$-crRNA-gapA targeting the *gapA* gene | This study |
| pcrRNA-zwf | pJYS2 derivative, $P_{j23119}$-crRNA-zwf targeting the *zwf* gene | This study |
| pcrRNA-pta | pJYS2 derivative, $P_{j23119}$-crRNA-pta targeting the *pta* gene | This study |
| pcrRNA-ackA | pJYS2 derivative, $P_{j23119}$-crRNA-ackA targeting the *ackA* gene | This study |
| pcrRNA-actA | pJYS2 derivative, $P_{j23119}$-crRNA-actA targeting the *actA* gene | This study |

**Supplementary Table S2.** Oligonucleotides used for gene cloning in this study.

| Name | Relevant characteristics (5’ → 3’) | Source |
|---|---|---|
| crRNA-*SpeI*-fwd | tatactagtATTTAAATAAAACGAAAGGC | This study |
| crRNA-*pfkA*-*SpeI*-rev | tatactagtCTTGATAACCAACGACGGTGGAGCATCTACAACAGTAGAAATTC | This study |
| crRNA-*zwf*-*SpeI*- rev | tatactagtAGCAATCCGCGGTTTGCTagATCAATCTACAACAGTAGAAATTC | This study |
| crRNA-*gapA*-*SpeI*- rev | tatactagtGCCGTATCggACGTAACTTCTTCCATCTACAACAGTAGAAATTC | This study |
| crRNA-*pta*-*SpeI*- rev | tatactagtCCAAGGTCTGCTGCTACTTCTTCATCTACAACAGTAGAAATTC | This study |
| crRNA-*actA*-*SpeI*- rev | tatactagtACCAACCTTGTCACCGTGGTTAAATCTACAACAGTAGAAATTC | This study |
| crRNA-*ackA*-*SpeI*- rev | tatactagtTTGATGGAAGATGAACCGGAGTTATCTACAACAGTAGAAATTC | This study |

Note: The restriction enzyme sites were shown as lower cases. Target specific-protospacer regions of the sgRNAs were underlined.

**Supplementary Table S3.** Single strand oligonucleotides used for the CoryneCR12-CDS_del

| Name | Relevant characteristics (5’ → 3’) | Source |
|---|---|---|
| ssODN_*pfkA*_100del | TCGCCTAACAGTCCTTCCCAACCGTCTTGATAACCAACGAGTCTTCCA TATTAAACCCATCACAACACCCGCGCCGGAAC | This study |
| ssODN_*zwf*_100del | GCACTTGCGGCATCGCGTACGTATTTTTCAAAGTCTTCTTCAAGTCGC CAGTGACACCGAAGATCACCATGCCGGAAGGG | This study |
| ssODN_*gap*_100del | CTTCGATGTGAGCCTTAGCCGCGTTTGCATCGGTGAAGAAATGGAGTC ATCGTCGTATTCAACTTCCTGGCCAAGGCGGC | This study |
| ssODN_*pta*_100del | AGAACTTTGGATACTTCTAGTTCTTCGTCGGGCAGGTAGGACATCGCC TTTCTAATTTCAGCCTGAACCTTCTCATTGAT | This study |
| ssODN_*actA*_100del | GTTGCCTGCACCGTGTGCTTCTTTAGCCCGGTTAGCGATTGCTTGGAG CGCAGCTTTTCTGAAGCAATGCGATCAGACAT | This study |
| ssODN_*ackA*_100del | ATTTTGAGTACGATGCGGCCGTTTGGCTCACCAATCTGCTTGCCATTA GCTGCGTCCTCCTGCCTGAATTGCTGTGATGG | This study |

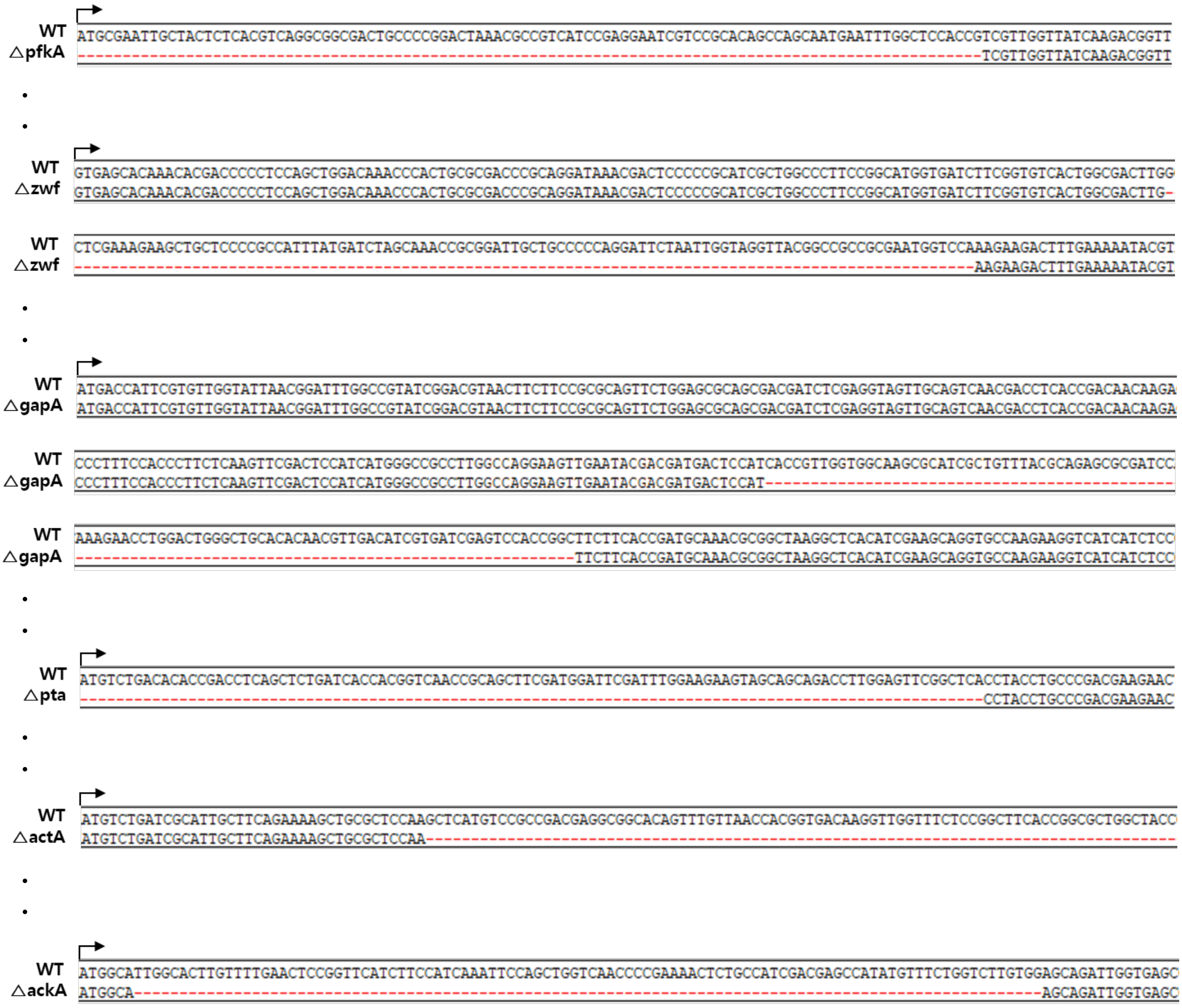


**Supplementary Fig. S1**. Sequence of editing sites at the *pfkA, zwf, gapA, pta, actA, ackA* gene of *C. glutamicum* constructed in this study. See the details in materials and methods for strain construction.

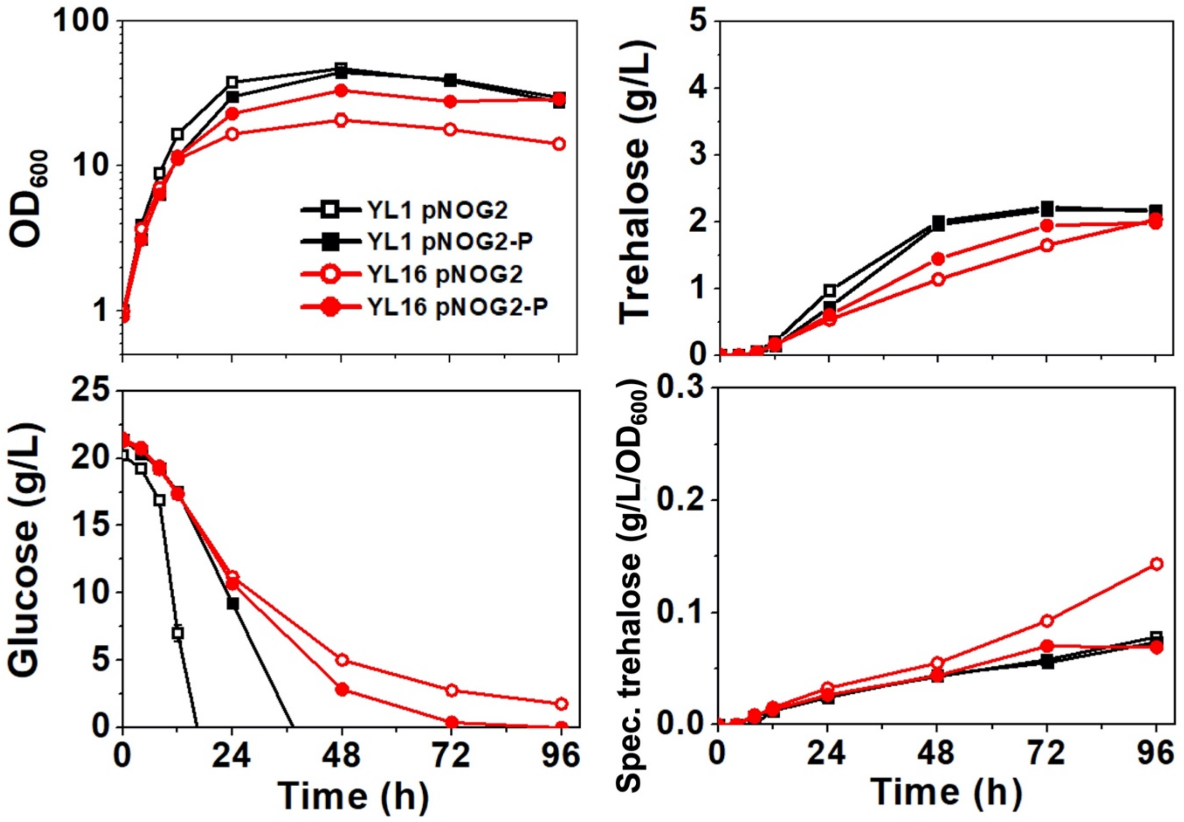


**Supplementary Fig. S2**. **Cell growth of *C. glutamicum* hybrid EMP variants.** Growth (optical density at 600 nm), glucose consumption (g/L), trehalose secretion (g/L), specific trehalose production (g/L/$OD_{600}$). YL1 pNOG2 was cultured in CgXII medium (50 mL in 250 mL baffled Erlenmeyer flasks) with 2% (w/v) glucose as the sole carbon source and various inorganic phosphate sources (13 mM as control, blue; 0.13 mM, black; 1.3 mM, red; 130 mM, green). Data represent mean values of at least triplicated cultivations, and error bars represent the standard deviations.

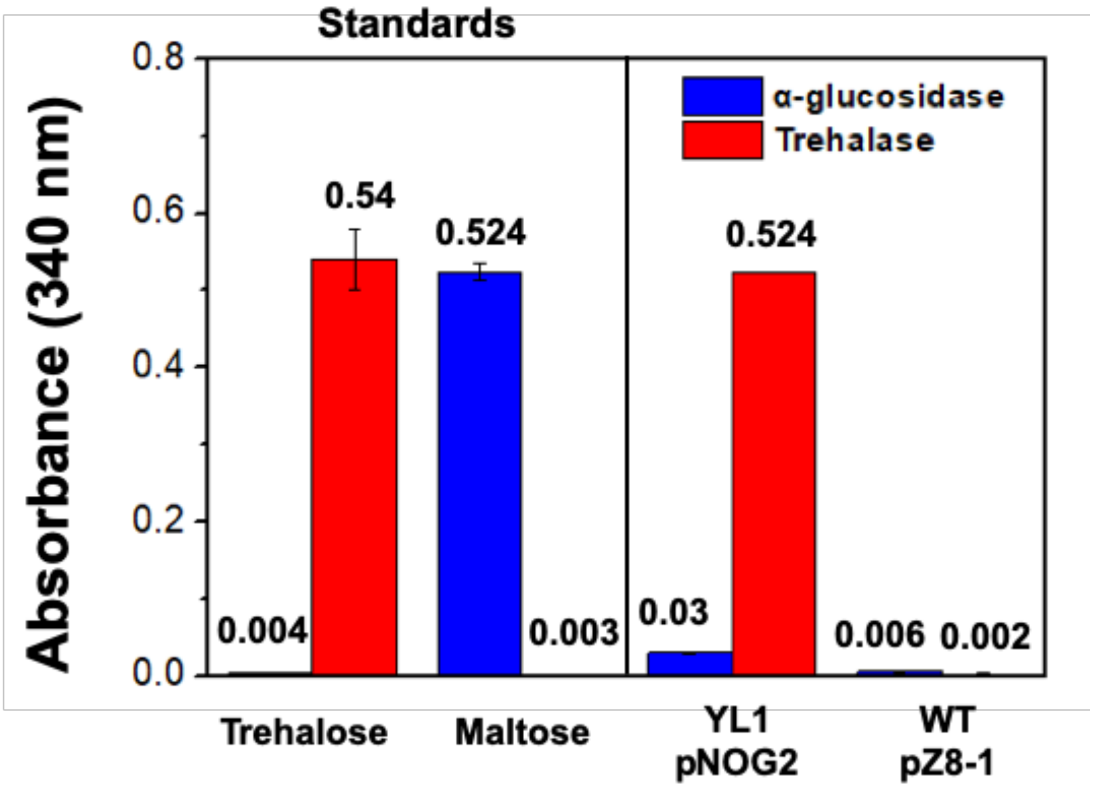


**Supplementary Fig. S3**. **Identification of trehalose secretion using the enzymatic assay**. Growth (optical density at 600 nm), glucose consumption (g/L), trehalose secretion (g/L), specific trehalose production (g/L/$OD_{600}$). YL1 pNOG2 was cultured in CgXII medium (50 mL in 250 mL baffled Erlenmeyer flasks) with 2% (w/v) glucose as the sole carbon source. WT pZ8-1 was used as a control strain. Trehalase (red bar) and α-glucosidase (blue bar) were used for enzymatic assay with diluted supernatants from either YL1 pNOG2 or WT pZ8-1, according to the range of the concentrations calculated using HPLC. Authentic trehalose and maltose standards were used as a control. Hexokinase and D-glulcose-6 phosphate dehydrogenase were coupled to detect free D-glucose by measuring the increase in the NAPDH levels at 340 nm using the kinetics module of the Eppendorf BioSpectrometer® (Eppendorf AG, Hamburg, Germany). All the procedures were performed according to the manufacturer's instructions. Standard chemicals (40 μg trehalose and 40 μg maltose; Sigma Aldrich) were also used as control during the enzymatic assays. Data represent mean values of at least triplicated cultivations, and error bars represent the standard deviations.
